# Qualitative Observations of Transgenically Individualized Serotonergic Fibers in the Mouse Telencephalon

**DOI:** 10.1101/2023.11.25.568688

**Authors:** Justin H. Haiman, Kasie C. Mays, Helen A. Roostaeyan, Nasim Elyasi, Skirmantas Janušonis

**Affiliations:** Department of Psychological and Brain Sciences, University of California, Santa Barbara, CA 93106-9660, USA

**Keywords:** 5-hydroxytryptamine (5-HT), serotonin, axon, fibers, densities, tortuosity, varicosities, Brainbow

## Abstract

The matrix of serotonergic axons (fibers) is a constant feature of neural tissue in vertebrate brains. Its fundamental role appears to be associated with the spatiotemporal control of neuroplasticity. The densities of serotonergic fibers vary across brain regions, but their development and maintenance remain poorly understood. A specific fiber concentration is achieved as the result of the dynamics of a large number of individual fibers, each of which can make trajectory decisions independently of other fibers. Bridging these processes, operating on very different spatial scales, remains a challenge in neuroscience. The study provides the first qualitative description of individually-tagged serotonergic axons in four selected telencephalic regions (cortical and subcortical) of the mouse brain. Based on our previous implementation of the Brainbow toolbox in this system, serotonergic fibers were labeled with random intensity combinations of three fluorophores and imaged with high-resolution confocal microscopy. All examined regions contained serotonergic fibers of diverse identities and morphologies, often traveling in close proximity to one another. Some fibers transitioned among several morphologies in the same imaged volume. High fiber densities appeared to be associated with highly tortuous fiber segments produced by some individual fibers. This study supports efforts to predictively model the self-organization of the serotonergic matrix in all vertebrates, including regenerative processes in the adult human brain.

## INTRODUCTION

All vertebrate brains contain a massive meshwork of serotonergic axons (fibers) that is produced by neurons located in the brainstem raphe nuclei, with a possible contribution by other nuclei (in non-mammals). These axons participate in neurosignaling by releasing 5-hydroxytryptamine (5-HT, serotonin) and can also release other neurotransmitters.^1^ The cellular origin and distribution of serotonergic fibers have been described in all vertebrate groups: the cartilaginous and bony fishes,^2-4^ the amphibians,^5^ the reptiles,^6^ the birds,^7,8^ and the mammals.^9,10^ The ubiquitous presence and high energetic cost of this fiber matrix suggests that it provides fundamental support for neural networks of the central nervous system. The current conceptualizations of this role remain grossly incomplete. The evolutionary forces that have shaped it likely predate the Permian-Triassic bottleneck 252 million years ago; the overarching functionality of the serotonergic matrix appears to be associated with neuroplasticity.^11-13^ The serotonergic matrix may maintain a dynamic balance between synaptic pliability and stability (metaphorically, keeping brain networks near the “melting temperature”). In particular, it is known to support transitions among global brain states such as wakefulness and sleep.^14^

The regional densities of serotonergic fibers have been described in many neuroanatomical studies,^15^ but their self-organization remains an unsolved problem. A number of signaling pathways appear to contribute to this process; the reported factors include Lmx1b,^16^ protocadherin-αC2,^17-19^ neurexins,^20^ S100β,^21^ the brain-derived neurotrophic factor (BDNF),^22^ and serotonin.^23,24^ Some aspects of this self-organization can be captured by models that treat the trajectories of serotonergic fibers as paths of stochastic processes.^15,25,26^ Ultimately, a comprehensive mechanistic explanation should bridge the dynamics of single serotonergic fibers – as they extend, branch, and react to chemical and mechanical signals – and their resultant densities in the adult brain. These densities may represent a dynamic equilibrium rather than a static distribution: serotonergic fibers can robustly regenerate in the adult mammalian brain, with new paths,^27,28^ which suggests that some individual fiber segments may continue to grow in the normal (injury-free) brain, with the accompanying degeneration of other segments.

Direct observations of single serotonergic fibers in brain tissue can support insights that are independent of current theoretical frameworks. Such observations not only complement hypothesis-driven research but are essential in inspiring new insights and intuitions that cannot be produced by logical inferences. A key challenge is this effort is that serotonergic fibers are thin, with a diameter that is near the limit of optical resolution, and that typically many fibers intertwine in the same three-dimensional volume. *In vitro* studies of isolated serotonergic neurons can provide some essential information,^29^ but cell-culture environments are radically different from those in the natural brain parenchyma. It should also be noted that analyses of the transcriptomes of individual serotonergic neurons have revealed an enormous diversity of transcriptional programs,^1,30,31^ further highlighting limitations of population-level studies.

We have recently developed a transgenic approach in which individual serotonergic fibers express random intensity combinations of three fluorophores with different spectral characteristics.^26^ This approach is an implementation of the Brainbow 3.2 toolbox,^32^ which is uniquely well-suited for this purpose. In particular, it allows for the identification of fiber segments that belong to the same or different neurons – even if these segments intersect, intertwine, or temporarily leave the imaging volume. This study is the first description of single serotonergic fibers with this level of resolution in a set of telencephalic regions of the mouse brain.

## RESULTS AND DISCUSSION

The three Brainbow fluorophores (EYFP, mTFP, TagBFP) were strongly expressed in the rostral raphe nuclei (Fig. 1), with no labeled somata outside the expected neuroanatomical region. In serotonergic axons, the mTFP signal (pseudocolored blue) had the lowest signal-to-noise ratio compared to the other two signals, but it contributed sufficiently to the diversification of the color signatures of individual fibers. In the following descriptions, fibers with “different identities” should be understood as fibers originating from different somata in the raphe complex. Generally, demonstrating that two fiber segments have different identities is easier than the opposite (since the intensities of the fluorophores are randomized in each neuron, some neurons may have similar color signatures).

**Figure 1.**
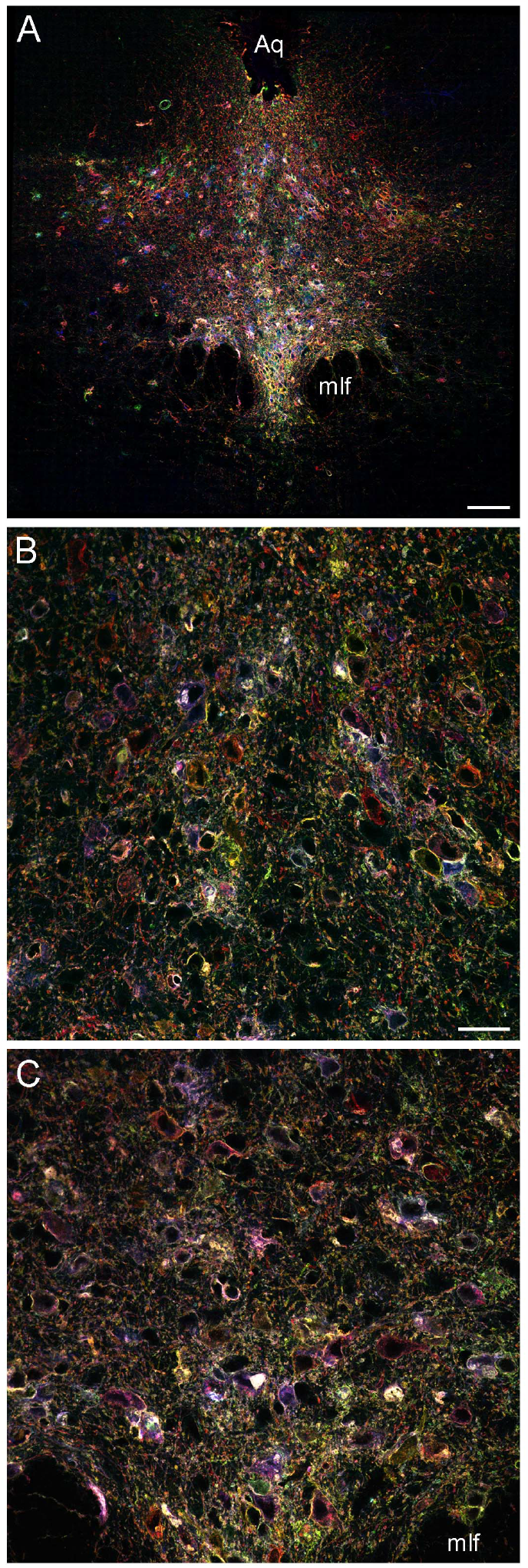
Confocal images of the dorsal raphe nucleus (DR). (**A**) One optical section of the entire nucleus (serotonergic neurons express different fluorophore intensity combinations; non-serotonergic neurons are not labeled). (**B**) One optical section of the dorsomedial DR. (**C**) One optical section of the ventromedial DR. Aq, aqueduct; mlf, medial longitudinal fasciculus. Scale bars = 100 μm in (A) and 30 μm in (B, C).

We examined four telencephalic regions: the primary somatosensory cortex (S1), the caudate-putamen nucleus, the immediate neighborhood of the anterior part of the anterior commissure, and the stratum lacunosum moleculare of the hippocampus. These regions represent major subdivisions of the mammalian telencephalon: the neocortex, the dorsal striatum, the ventral striatum (with the extended amygdala), and the archicortex, respectively. Serotonergic fibers can travel with no large physical obstacles in the neocortex and caudate-putamen (with the exception of the ventricular and pial boundaries). In contrast, the anterior commissure, an ancient interhemispheric tract that phylogenetically precedes the corpus callosum,^33^ is virtually impenetrable to serotonergic fibers and therefore a major obstacle located inside neural tissue.

Not all neuroanatomical tracts have this property: for example, serotonergic fibers can partially penetrate the fasciculus retroflexus^34,35^ and many of them are present in the stria terminalis.^9^ The stratum lacunosum moleculare has one of the highest serotonergic fiber densities in the entire brain^9^; these fibers can be affected by selective serotonin reuptake inhibitors (SSRIs).^24^

In the somatosensory cortex, the density of serotonergic fibers progressed from a relatively high density in layer 1 to a relatively sparse distribution in layers 2-3 (Fig. 2A), consistent with previous descriptions that have used monochrome visualization methods.^9,34^ In these low-density layers, most fibers had low tortuosity and some fibers branched frequently (Figs. 2B-D). Limited information can be extracted from fixed tissue, but these high-resolution images suggest that fiber branches can develop by lateral “budding” (in which one branch is developmentally older than the other one) and by bifurcation (where both branches are of the same age). Both mechanisms have been proposed in previous studies,^36-38^ but this question can be definitely answered only with modern live imaging methods (e.g., 3D-holotomography^29^). In layer 1, fibers of distinctly different identities traveled together (separated by only several micrometers), with no detectable interference (Fig. 2E). Some of these fibers also strongly differed in their morphologies, ranging from those with strong varicosities and virtually undetectable intervaricose segments to those with a smooth or “undulating” appearance (Figs. 2F-J). Within the small imaged volume (around 10^6^ μm^3^), individual fibers tended to maintain the same morphology. However, some longer segments underwent transitions from relatively straight, thin segments to dilated segments (Fig. 2K). Our previous *in vitro* analysis has demonstrated similar transitions *in vitro*, also with two branches of the same growing serotonergic axon differing considerably in their morphological appearance.^29^

**Figure 2.**
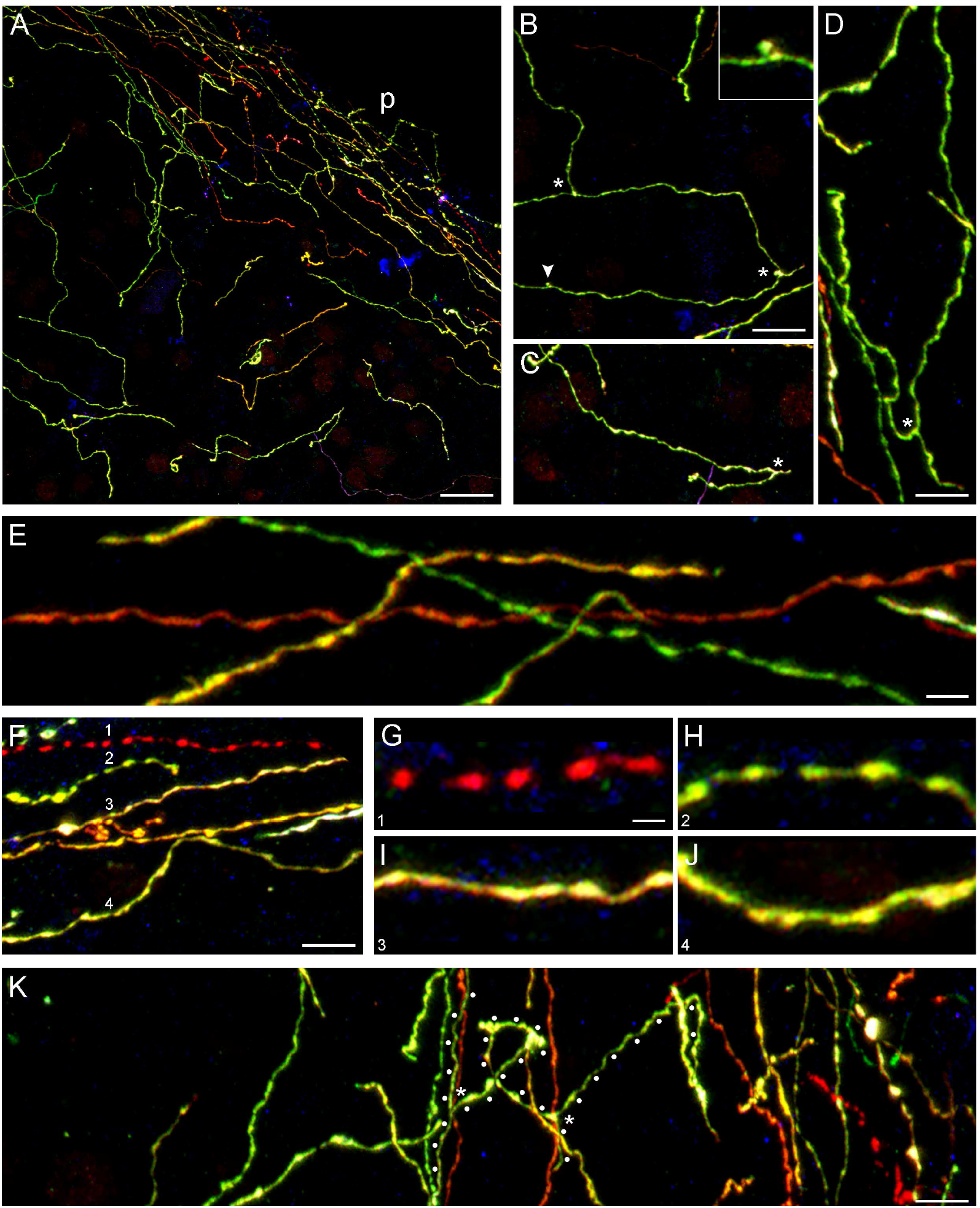
(**A**) A confocal image of layers 1-3 of the primary somatosensory cortex (the barrel field region). All remaining panels are subsets of it (some with rotation to optimize the layout). The pial border (p) is shown. (**B-D**) Frequent branching events in fiber segments that have similar color signatures (and may be the same fiber). An inset (B) shows a potential budding event that might produce a new branching point (4× magnification). (**E**) Several fibers with distinctly different color signatures that travel together. (**F**) Several fibers with different color signatures that travel together but show distinctly different morphologies (shown in detail in (G-J), with matching numbers). Fiber 1 (red) is closest to the pia. (**G**) A fiber with clear varicosities and virtually undetectable intervaricose segments. (**H**) A fiber with clear varicosities and thin but detectable intervaricose segments. (**I**) A relatively smooth fiber, with no pronounced varicosities. (**J**) A fiber that may have active membrane expansions similar to those observed in growing axons.^29,36^ (**K**) A fiber that transitions between smooth, relatively straight segments and tortuous segments of a larger caliber. The fiber path is marked with white dots and the branching points with asterisks. Scale bars = 20 μm in (A), 10 μm in (B, C), 5 μm in (D, F, K), 2 μm in (E), and 1 μm in (G-J).

The caudate-putamen and the bed nucleus of the stria terminalis (BNST) are adjacent at some coronal levels (Fig. 3A) but differ in both their serotonergic fiber densities and their position with respect to large physical obstacles. Axon fascicles are abundant in the caudate-putamen^39^ but do not appear to significantly hinder the movement of serotonergic fibers (Fig. 3B). In contrast, serotonergic fibers are physically constrained in some subregions of the BNST by the lateral ventricle and the anterior commissure (Fig. 3C). The caudate-putamen had a moderate density of serotonergic fibers (Fig. 4A), consistent with a published monochrome map.^9^ As in the primary somatosensory cortex, many fibers in the caudate-putamen maintained the same direction (Fig. 4B), fibers of different identities and morphologies traveled together (Fig. 4C), and some fibers showed reversible transitions between different morphologies (Fig. 3D, E). The BNST had a high density of serotonergic fibers (Fig. 4F), in register with the monochrome map.^9^ This higher condensation of fibers was driven, at least in part, by islands of highly intertwined, tortuous fiber segments (Fig. 4G). These tangles had no discernable organization; many were produced by fibers of clearly different identities (Fig. 4H).

**Figure 3.**
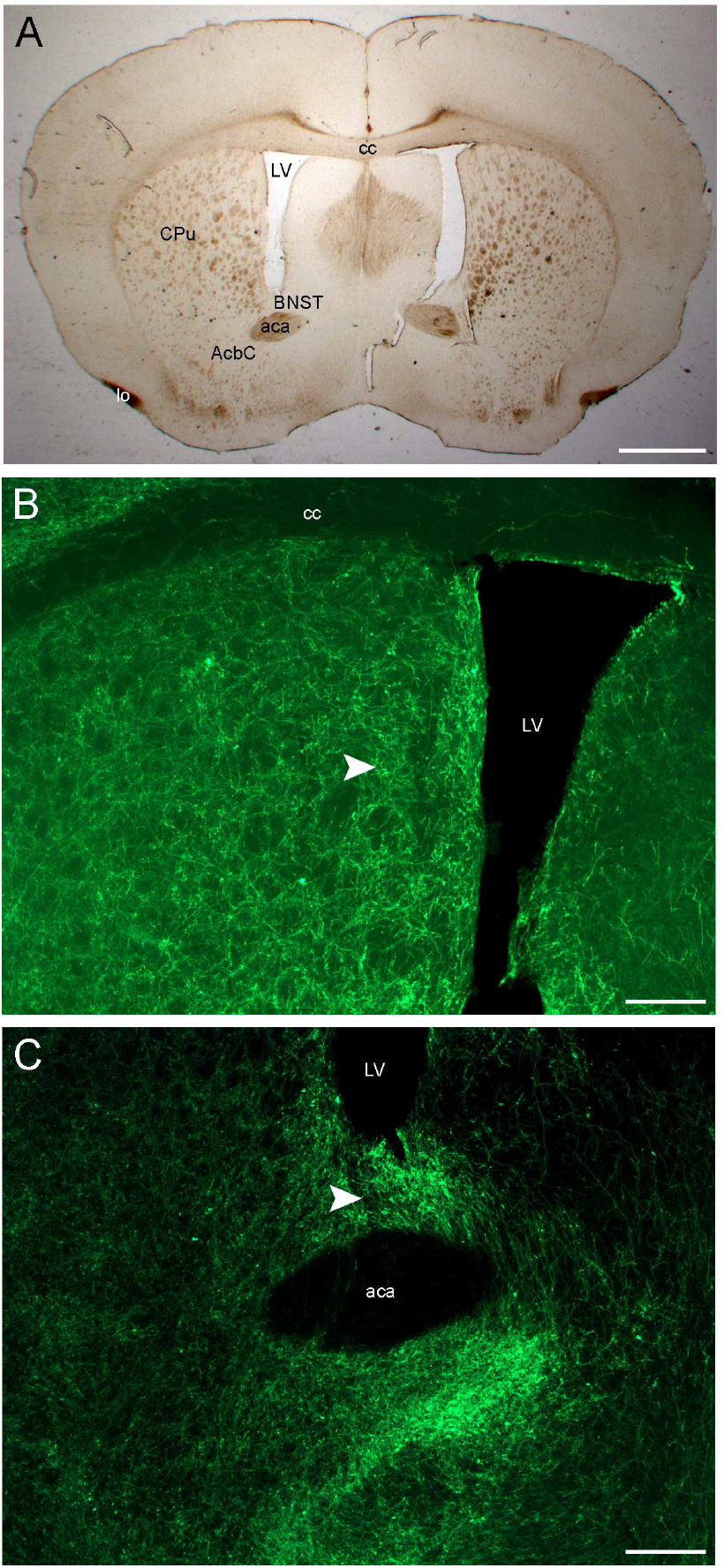
(**A**) A low-power, bright-field image of the section used to capture the confocal images in Figure 4. The AlexaFluor dyes are not visible; no additional staining was used. (**B**) An epifluorescent image (in the AlexaFluor 488 channel) of the core of the caudate putamen. The arrowhead shows the region imaged with confocal microscopy. (**C**) An epifluorescent image (in the AlexaFluor 488 channel) of a high-density region between the lateral ventricle and the anterior commissure. The arrowhead shows the region imaged with confocal microscopy. aca, anterior commissure (anterior part); AcbC, the nucleus accumbens (core); BNST, bed nucleus of the stria terminalis; cc, corpus callosum; CPu, caudate putamen; LV, lateral ventricle. Scale bars = 1 mm in (A), 200 μm in (B, C).

**Figure 4.**
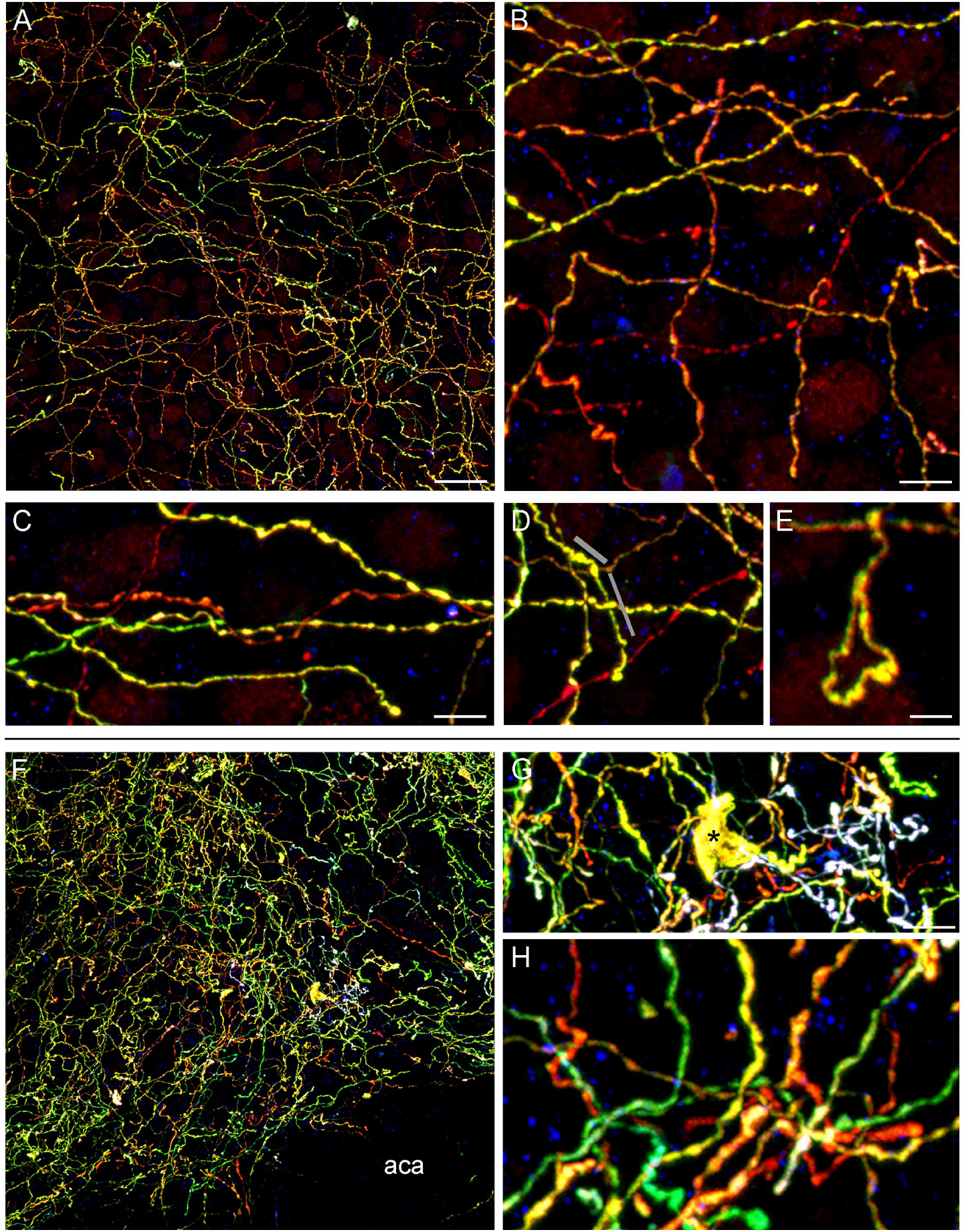
(**A**) A confocal image of the caudate-putamen. Panels (B-E) are subsets of it (some with rotation to optimize the layout). (**B**) Fibers with low tortuosity that create a relatively sparse meshwork. (**C**) Several fibers that travel together but show distinctly different morphologies. (**D**) A fiber that changes its morphology from a strongly dilated segment to a smooth segment (marked with gray bars). (**E**) A loop configuration. (**F**) A confocal image of the core of the nucleus accumbens at the same coronal level. The anterior commissure (the anterior part, aca) is virtually impenetrable. Panels (G, H) are subsets of this image (with no rotation). (**G**) Tortuous and densely packed fibers near the anterior commissure. The large structure marked with an asterisk is unidentified but does not appear to be an artifact; similar structures are present in other locations of the confocal image. (**H**) Fibers of several identities intertwine to produce high-density clusters. Scale bars = 20 μm in (A), 5 μm in (B), 5 μm in (C, D), 2 μm in (E), 20 μm in (F), 5 μm in (G), and 2 μm in (H).

The stratum lacunosum moleculare had a high density of serotonergic fibers (Fig. 5A), consistent with the monochrome map.^9^ Some fibers produced relatively straight trajectories, apparently encountering no obstacles for distances equivalent of several soma diameters (Fig. 5B). The high fiber density was supported by multiple, scattered islands of highly tortuous fiber segments (Fig. 5C). Interestingly, many of these islands had the same color signature (also accompanied by comparable fiber morphologies), suggesting that they were produced by the same fiber (Figs. 5D-F). If this conclusion is correct, high serotonergic fiber densities in brain regions can be built by fibers that switch between low and high tortuosity regimes, resulting in sparse and concentrated microregions, respectively (Fig. 5D). We have observed similar profiles in the mouse habenula.^26^ Theoretical tools have been recently developed to model this and similar dynamics.^25,26^ The high-tortuosity mode may be a property intrinsic to the fiber, but it may also be induced by its environment. For example, it may be associated with pericellular baskets that serotonergic neurons fibers can produce.^40^

**Figure 5.**
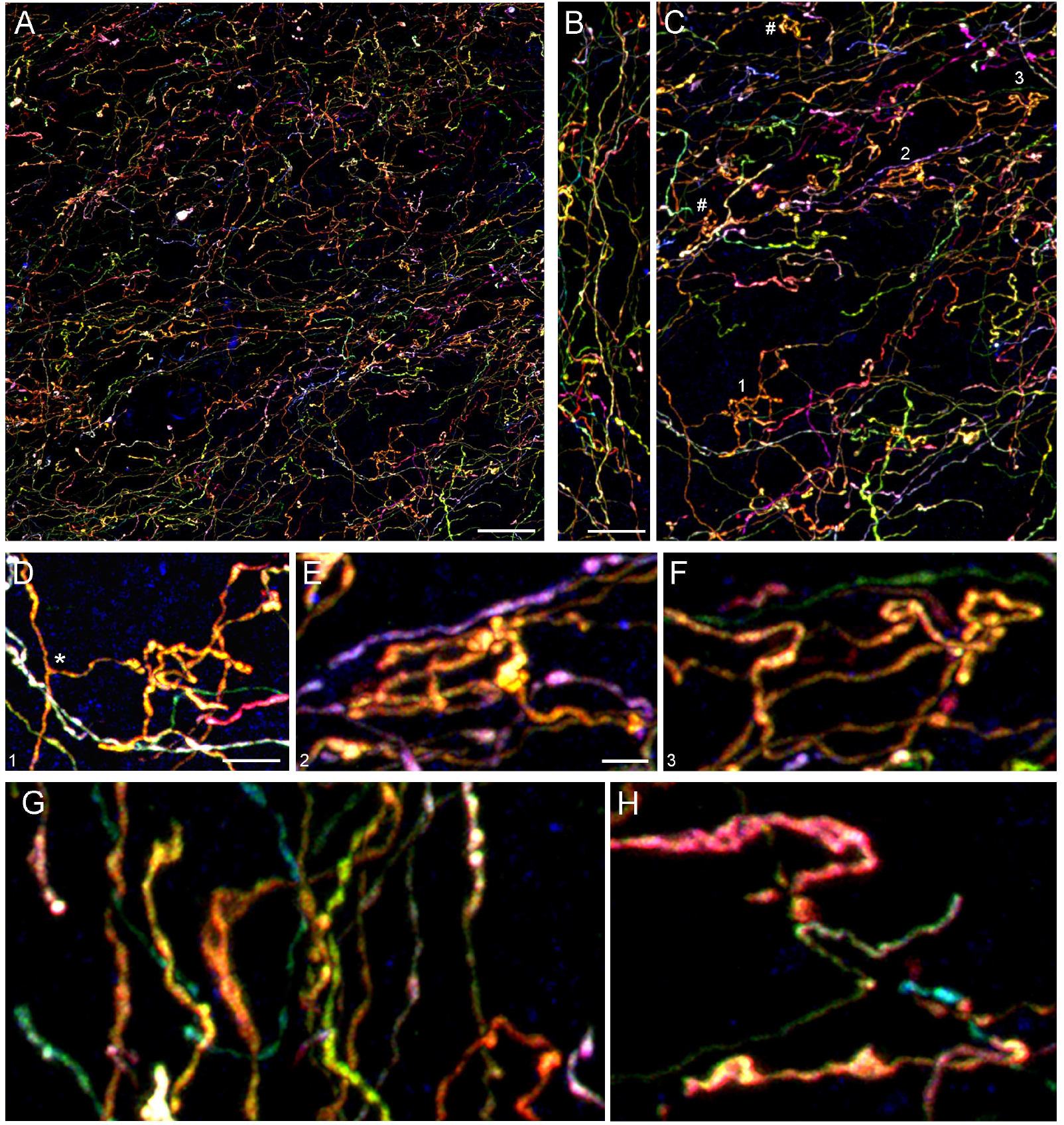
(**A**) A confocal image of the lacunosum moleculare of the CA1 field of the hippocampus. All remaining panels are subsets of it (some with rotation to optimize the layout). (**B**) A fiber that travels straight for over 60 μm. (**C**) Highly tortuous islands produced in different locations by fiber segments that have the same color signature (orange) and are likely to belong to the same fiber. The islands are labeled with numbers (1-3) and two hash signs (#). (**D-F**) High-power images of the numbered islands. (**G**) Several fibers with different color signatures that travel together but show distinctly different morphologies. The dilated appearance of one fiber (arrowhead) may be due to its flattened morphology. (**H**) Oher segments with a similar appearance. Scale bars = 20 μm in (A), 10 μm in (B, C), 5 μm in (D), and 2 μm in (E-H).

As in the other studied regions, fibers of multiple identities and morphologies often traveled together in the hippocampus (Fig. 5G). Some fiber segments appeared strongly dilated (Figs. 5G, H). These segments resembled profiles captured with superresolution microscopy in the embryonic mouse brain and with holotomography *in vitro* and may be flat.^29^

The observations in the selected regions suggest that serotonergic fibers of many identities are present in many, if not all, locations of neural tissue. These fibers also often differ in their morphologies (e.g., the presence and appearance of varicosities, as well as the caliber). Since their segments co-occupy the same small tissue volume, these differences are unlikely to be induced by local signals in the environment. They may instead reflect individually-specific transcriptome programs of the corresponding somata,^1,30,31^ the individual age of the segments,^41^ or their unique individual history (e.g., a certain appearance may be induced a physical obstacle resisting fiber extension and may persist for some time after the event). The latter two possibilities are supported by observations that serotonergic fibers can regenerate in the adult brain^27^ and that the current extension of a fiber may depend of the history of its previous extensions.^15,34^.

The selected region set is small and not fully representative. However, together with our recent introduction of this methodological approach,^26^ it presents the first high-resolution view of individually-tagged serotonergic fibers in the brain. This vast and elegant system will challenge commonly held beliefs and require more direct observations (analogous to field studies in biology) – before its rules and patterns have been well understood. On the biomedical front, this understanding will support efforts to preserve the health of neural tissue and to support its regeneration.^42^

## METHODS

### General Approach

We used an implementation of the Brainbow 3.2 transgenic toolbox^32^ in the brain serotonergic system, which has been recently developed in our laboratory.^26^ The method is described here briefly, with some further modifications.

### Animals

A transgenic mouse line that expresses a tamoxifen-inducible Cre recombinase under the control of the promoter of the *Tph2* gene (a rate-limiting enzyme in the serotonin synthesis pathway) was used. Founding breeders, which included mice on the same genetic background without the transgene, were purchased from The Jackson Laboratory (JAX). The breeding colony was kept in a vivarium on a 12:12 light-dark cycle with free access to food and water. Transgene-carriers were mated with non-carriers, and the litters were PCR-genotyped by using toe biopsies collected at postnatal day 7. All animal procedures have been approved by the UCSB Institutional Animal Care and Use Committee.

### Intracranial Injections of Brainbow AAVs

The Brainbow adeno-associated viruses (AAVs) (AAV-EF1a-BbTagBY [Addgene #45185-AAV9] and AAV-EF1a-BbChT [Addgene #45186-AAV9]) (Cai et al., 2013) were diluted and mixed before use to 1.5×10^12^ vg/mL each in a tube with sterile, alcohol-free saline. An adult mouse with the transgene was anesthetized with a mixture of ketamine (100 mg/kg) and xylazine (10 mg/kg), and further anesthesia was maintained with inhaled isoflurane. The animal was placed in a stereotaxic apparatus, and a small hole in the skull was drilled over the dorsal raphe (DR), after a skin incision and retraction. The AAVs (1-2 μL) were slowly pressure-injected intracranially with a 10 μL-Hamilton microsyringe into the DR (at lambda, 3.5 mm ventral to the dura). This volume is sufficient to cover the entire region that contains the rostral raphe nuclei (without diffusion, it is equivalent to a sphere with the diameter of 1.2-1.6 mm). The needle was kept in the DR for 5-10 minutes and then slowly withdrawn. The skin incision was closed with sterile wound clips, and the animal was monitored until full recovery. Four animals (all males) were used in the study. The study did not include any of the animals used in the previous study^26^ (all AAV injections were performed in 2023).

### Induction of Cre Recombination

One week after the surgery, the induction of Cre-recombination was achieved by administering tamoxifen (Millipore-Sigma #T5648) dissolved in corn oil (Millipore-Sigma #C8267). The stock concentration (20 mg/mL), stored at -20°C, was injected intraperitoneally (0.1 mL at 75 mg/kg) for 5 consecutive days. Following the treatment, the mice were allowed to survive for three months to ensure fluorophore transport to distal axon segments.

### Brainbow Immunohistochemistry

Mice were euthanized with CO_2_ and their brains were dissected into 0.1 M phosphate-buffered saline (PBS, pH 7.2). They were immediately immersion-fixed in 4% paraformaldehyde overnight at 4ºC, cryoprotected in 30% sucrose for two days at 4ºC, and sectioned coronally into PBS at 40 μm thickness on a freezing microtome. Sections through the rostral raphe were examined unstained with epifluorescence microscopy, and two mice (100 and 111 days post-surgery) with excellent fluorophore expression in the entire raphe region were selected for triple-label immunohistochemistry (with some modifications of the original protocol^26^). At the time of the brain dissection, the mice were 17 and 17.5 months of age, respectively. Meaningful age comparisons across mammalian clades are difficult; however, this age has been considered late middle adulthood (or roughly 50 human years).^43,44^ Sections were rinsed in PBS, blocked in 5% normal donkey serum (NDS) for 30 minutes at room temperature, and incubated on a shaker in rabbit anti-GFP IgG (1:500; Abcam #ab6556), rat anti-mTFP IgG (1:200; Ximbio #155264), and guinea pig anti-TagRFP IgG (1:500; Ximbio #155267), with 3% NDS and 0.5% TX in PBS for 3 days at 4°C. The anti-GFP and anti-TagRFP antibodies recognize EYFP and TagBFP, respectively. The sections were rinsed in PBS three times (10 minutes each) and incubated in AlexaFluor 488-conjugated donkey anti-rabbit IgG (1:500; ThermoFisher #A21206), AlexaFluor 594-conjugated donkey anti-rat IgG (1:500; Jackson ImmunoResearch #712-585-150), and AlexaFluor 647-conjugated donkey anti-guinea pig IgG (1:500; Jackson ImmunoResearch #706-605-148) with 3% NDS in PBS for 90 minutes at room temperature. The sections were rinsed in PBS three times (10 minutes each), mounted onto gelatin/chromium-subbed glass slides, allowed to air-dry, coverslipped with ProLong Gold Antifade Mountant with no DAPI (ThermoFisher Scientific, #P36930), and allowed to cure for over 24 hours.

### Microscopy Imaging

Bright-field and one-channel (GFP) epifluorescence images were captured on a Zeiss AxioVision Z1 system with a 1× (NA 0.025) or 10× (NA 0.45) objectives. Confocal imaging was performed in three Brainbow channels (AlexaFluor 488, AlexaFluor 594 or 546, and AlexaFluor 647) on a Leica SP8 resonant scanning confocal system with a 63× objective (oil immersion, NA 1.40), at 72 nm/pixel (the XY plane) and 299 nm/optical section (the Z-axis). An optimal balance of the intensities in the AlexaFluor 488, AlexaFluor 594, and AlexaFluor 647 channels was achieved with approximately 30%, 15%, and 4% laser power, respectively. These channels were pseudocolored green, blue, and red, respectively. The highest-intensity pixels were allowed to saturate, but the proportion of the saturated pixels was kept minimal and carefully balanced with the signal intensity, to minimize non-linearities and subsequent color shifts (e.g., if all three channels are saturated, the resultant color is always white). Confocal z-stacks contained around 100 optical sections (2552 × 2552 pixels). The extreme optical sections at the top and bottom of the stack were removed (to reduce edge artifacts), a Gaussian-blur filter with a kernel of 3 pixels was applied in each channel, the maximum-intensity projections was computed, and the pixel values were rescaled in each channel to utilize the entire dynamic range of the values (0-100% brightness). The confocal images in Figure 1 were captured with 10× (NA 0.30) and 40× (water immersion, NA 1.10) objectives and show single optical sections (4.3 μm and 0.4 μm thick, respectively). In descriptions, a “color” should be interpreted as a specific set of intensities in the three channels (each one of which contains grayscale pixel values).

## Author Contributions

J.H.H. led and performed all experiments based on the method developed by K.C.M. H.A.R and N.E. provided essential technical assistance. S.J. supervised the project, performed some confocal microscopy analyses, and wrote the manuscript.

## Notes

The authors declare no competing financial interest.

## ACKNOWLEDGEMENTS

This research was supported by the National Science Foundation grants #2112862 and the California NanoSystems Institute (Challenge-Program Development grants). We acknowledge the use of the NRI-MCDB Microscopy Facility and the Leica SP8 Resonant Scanning Confocal Microscope (supported by National Science Foundation MRI grant #1625770).

## Notes

### Competing Interest Statement

The authors have declared no competing interest.

## REFERENCES

(1) Okaty, B. W.; Commons, K. G.; Dymecki, S. M. Embracing diversity in the 5-HT neuronal system. Nat Rev Neurosci 2019, 20 (7), 397–424. DOI: 10.1038/s41583-019-0151-3

(2) Carrera, I.; Molist, P.; Anadon, R.; Rodriguez-Moldes, I. Development of the serotoninergic system in the central nervous system of a shark, the lesser spotted dogfish Scyliorhinus canicula. J Comp Neurol 2008, 511 (6), 804–831. DOI: 10.1002/cne.21857

(3) Janušonis, S. Some galeomorph sharks express a mammalian microglia-specific protein in radial ependymoglia of the telencephalon. Brain Behav Evol 2018, 91 (1), 17–30. DOI: 10.1159/000484196

(4) Lillesaar, C. The serotonergic system in fish. J Chem Neuroanat 2011, 41 (4), 294–308. DOI: 10.1016/j.jchemneu.2011.05.009

(5) Bhat, S. K.; Ganesh, C. B. Organization of serotonergic system in Sphaerotheca breviceps (Dicroglossidae) tadpole brain. Cell Tissue Res 2023, 391 (1), 67–86. DOI: 10.1007/s00441-022-03709-7

(6) Bennis, M.; Gamrani, H.; Geffard, M.; Calas, A.; Kah, O. The distribution of 5-HT immunoreactive systems in the brain of a saurian, the chameleon. J Hirnforsch 1990, 31 (5), 563–574.

(7) Metzger, M.; Toledo, C.; Braun, K. Serotonergic innervation of the telencephalon in the domestic chick. Brain Res Bull 2002, 57 (3-4), 547–551. DOI: 10.1016/s0361-9230(01)00688-8

(8) Metzger, M.; Britto, L. R.; Toledo, C. A. Monoaminergic markers in the optic tectum of the domestic chick. Neuroscience 2006, 141 (4), 1747–1760. DOI: 10.1016/j.neuroscience.2006.05.020

(9) Awasthi, J. R.; Tamada, K.; Overton, E. T. N.; Takumi, T. Comprehensive topographical map of the serotonergic fibers in the male mouse brain. J Comp Neurol 2021, 529 (7), 1391–1429. DOI: 10.1002/cne.25027 From NLM.

(10) Hornung, J. P. The human raphe nuclei and the serotonergic system. J Chem Neuroanat 2003, 26 (4), 331–343. DOI: 10.1016/j.jchemneu.2003.10.002

(11) Lesch, K. P.; Waider, J. Serotonin in the modulation of neural plasticity and networks: implications for neurodevelopmental disorders. Neuron 2012, 76 (1), 175–191. DOI: 10.1016/j.neuron.2012.09.013

(12) Morgan, A. A.; Alves, N. D.; Stevens, G. S.; Yeasmin, T. T.; Mackay, A.; Power, S.; Sargin, D.; Hanna, C.; Adib, A. L.; Ziolkowski-Blake, A.; et al. Medial prefrontal cortex serotonin input regulates cognitive flexibility in mice. bioRxiv 2023. DOI: 10.1101/2023.03.30.534775

(13) Girn, M.; Rosas, F. E.; Daws, R. E.; Gallen, C. L.; Gazzaley, A.; Carhart-Harris, R. L. A complex systems perspective on psychedelic brain action. Trends Cogn Sci 2023, 27 (5), 433–445. DOI: 10.1016/j.tics.2023.01.003

(14) Brown, R. E.; Basheer, R.; McKenna, J. T.; Strecker, R. E.; McCarley, R. W. Control of sleep and wakefulness. Physiol Rev 2012, 92 (3), 1087–1187. DOI: 10.1152/physrev.00032.2011

(15) Janušonis, S.; Haiman, J. H.; Metzler, R.; Vojta, T. Predicting the distribution of serotonergic axons: a supercomputing simulation of reflected fractional Brownian motion in a 3D-mouse brain model. Front Comput Neurosci 2023, 17, 1189853. DOI: 10.3389/fncom.2023.1189853

(16) Donovan, L. J.; Spencer, W. C.; Kitt, M. M.; Eastman, B. A.; Lobur, K. J.; Jiao, K.; Silver, J.; Deneris, E. S. Lmx1b is required at multiple stages to build expansive serotonergic axon architectures. Elife 2019, 8, e48788. DOI: 10.7554/eLife.48788

(17) Katori, S.; Hamada, S.; Noguchi, Y.; Fukuda, E.; Yamamoto, T.; Yamamoto, H.; Hasegawa, S.; Yagi, T. Protocadherin-alpha family is required for serotonergic projections to appropriately innervate target brain areas. J Neurosci 2009, 29 (29), 9137–9147. DOI: 10.1523/jneurosci.5478-08.2009

(18) Katori, S.; Noguchi-Katori, Y.; Okayama, A.; Kawamura, Y.; Luo, W.; Sakimura, K.; Hirabayashi, T.; Iwasato, T.; Yagi, T. Protocadherin-αC2 is required for diffuse projections of serotonergic axons. Sci Rep 2017, 7 (1), 15908. DOI: 10.1038/s41598-017-16120-y

(19) Chen, W. V.; Nwakeze, C. L.; Denny, C. A.; O’Keeffe, S.; Rieger, M. A.; Mountoufaris, G.; Kirner, A.; Dougherty, J. D.; Hen, R.; Wu, Q.; et al. Pcdhaαc2 is required for axonal tiling and assembly of serotonergic circuitries in mice. Science 2017, 356 (6336), 406–411. DOI: 10.1126/science.aal3231

(20) Cheung, A.; Konno, K.; Imamura, Y.; Matsui, A.; Abe, M.; Sakimura, K.; Sasaoka, T.; Uemura, T.; Watanabe, M.; Futai, K. Neurexins in serotonergic neurons regulate neuronal survival, serotonin transmission, and complex mouse behaviors. Elife 2023, 12. DOI: 10.7554/eLife.85058

(21) Whitaker-Azmitia, P. M. Serotonin and brain development: role in human developmental diseases. Brain Res Bull 2001, 56 (5), 479–485. DOI: 10.1016/s0361-9230(01)00615-3

(22) Mamounas, L. A.; Blue, M. E.; Siuciak, J. A.; Altar, C. A. Brain-derived neurotrophic factor promotes the survival and sprouting of serotonergic axons in rat brain. J Neurosci 1995, 15 (12), 7929–7939.

(23) Migliarini, S.; Pacini, G.; Pelosi, B.; Lunardi, G.; Pasqualetti, M. Lack of brain serotonin affects postnatal development and serotonergic neuronal circuitry formation. Mol Psychiatry 2013, 18 (10), 1106–1118. DOI: 10.1038/mp.2012.128

(24) Nazzi, S.; Maddaloni, G.; Pratelli, M.; Pasqualetti, M. Fluoxetine induces morphological rearrangements of serotonergic fibers in the hippocampus. ACS Chem Neurosci 2019, 10 (7), 3218–3224. DOI: 10.1021/acschemneuro.8b00655

(25) Janušonis, S.; Detering, N. A stochastic approach to serotonergic fibers in mental disorders. Biochimie 2019, 161, 15–22. DOI: 10.1016/j.biochi.2018.07.014

(26) Mays, K. C.; Haiman, J. H.; Janušonis, S. An experimental platform for stochastic analyses of single serotonergic fibers in the mouse brain. Front Neurosci 2023, 17, 1241919. DOI: 10.3389/fnins.2023.1241919

(27) Jin, Y.; Dougherty, S. E.; Wood, K.; Sun, L.; Cudmore, R. H.; Abdalla, A.; Kannan, G.; Pletnikov, M.; Hashemi, P.; Linden, D. J. Regrowth of serotonin axons in the adult mouse brain following injury. Neuron 2016, 91 (4), 748–762. DOI: 10.1016/j.neuron.2016.07.024

(28) Cooke, P.; Janowitz, H.; Dougherty, S. E. Neuronal redevelopment and the regeneration of neuromodulatory axons in the adult mammalian central nervous system. Front Cell Neurosci 2022, 16, 872501. DOI: 10.3389/fncel.2022.872501

(29) Hingorani, M.; Viviani, A. M. L.; Sanfilippo, J. E.; Janušonis, S. High-resolution spatiotemporal analysis of single serotonergic axons in an in vitro system. Front Neurosci 2022, 16, 994735. DOI: 10.3389/fnins.2022.994735

(30) Okaty, B. W.; Sturrock, N.; Escobedo Lozoya, Y.; Chang, Y.; Senft, R. A.; Lyon, K. A.; Alekseyenko, O. V.; Dymecki, S. M. A single-cell transcriptomic and anatomic atlas of mouse dorsal raphe Pet1 neurons. Elife 2020, 9. DOI: 10.7554/eLife.55523

(31) Ren, J.; Isakova, A.; Friedmann, D.; Zeng, J.; Grutzner, S. M.; Pun, A.; Zhao, G. Q.; Kolluru, S. S.; Wang, R.; Lin, R.; et al. Single-cell transcriptomes and whole-brain projections of serotonin neurons in the mouse dorsal and median raphe nuclei. Elife 2019, 8. DOI: 10.7554/eLife.49424

(32) Cai, D.; Cohen, K. B.; Luo, T.; Lichtman, J. W.; Sanes, J. R. Improved tools for the Brainbow toolbox. Nat Methods 2013, 10 (6), 540–547

(33) Fenlon, L. R.; Suarez, R.; Lynton, Z.; Richards, L. J. The evolution, formation and connectivity of the anterior commissure. Semin Cell Dev Biol 2021, 118, 50–59. DOI: 10.1016/j.semcdb.2021.04.009

(34) Janušonis, S.; Detering, N.; Metzler, R.; Vojta, T. Serotonergic axons as fractional Brownian motion paths: Insights into the self-organization of regional densities. Front Comput Neurosci 2020, 14, 56. DOI: 10.3389/fncom.2020.00056

(35) Lidov, H. G.; Molliver, M. E. An immunohistochemical study of serotonin neuron development in the rat: ascending pathways and terminal fields. Brain Res Bull 1982, 8 (4), 389–430. DOI: 10.1016/0361-9230(82)90077-6

(36) Bovolenta, P.; Mason, C. Growth cone morphology varies with position in the developing mouse visual pathway from retina to first targets. J Neurosci 1987, 7 (5), 1447–1460. DOI: 10.1523/jneurosci.07-05-01447.1987

(37) Norris, C. R.; Kalil, K. Morphology and cellular interactions of growth cones in the developing corpus callosum. J Comp Neurol 1990, 293 (2), 268–281. DOI: 10.1002/cne.902930209

(38) Gallo, G.; Letourneau, P. C. Regulation of growth cone actin filaments by guidance cues. J Neurobiol 2004, 58 (1), 92–102. DOI: 10.1002/neu.10282

(39) Ni, R. J.; Huang, Z. H.; Shu, Y. M.; Wang, Y.; Li, T.; Zhou, J. N. Atlas of the striatum and globus pallidus in the tree shrew: Comparison with rat and mouse. Neurosci Bull 2018, 34 (3), 405–418. DOI: 10.1007/s12264-018-0212-z

(40) Senft, R. A.; Dymecki, S. M. Neuronal pericellular baskets: neurotransmitter convergence and regulation of network excitability. Trends Neurosci 2021, 44 (11), 915–924. DOI: 10.1016/j.tins.2021.08.006

(41) Maddaloni, G.; Bertero, A.; Pratelli, M.; Barsotti, N.; Boonstra, A.; Giorgi, A.; Migliarini, S.; Pasqualetti, M. Development of serotonergic fibers in the post-natal mouse brain. Front Cell Neurosci 2017, 11, 202. DOI: 10.3389/fncel.2017.00202

(42) Bradke, F. Mechanisms of Axon growth and regeneration: moving between development and disease. J Neurosci 2022, 42 (45), 8393–8405. DOI: 10.1523/jneurosci.1131-22.2022

(43) Hagan, C. When are mice considered old? The Jackson Laboratory 2017 (November 7, 2017)

(44) McWain, M. A.; Pace, R. L.; Nalan, P. A.; Lester, D. B. Age-dependent effects of social isolation on mesolimbic dopamine release. Exp Brain Res 2022, 240 (10), 2803–2815. DOI: 10.1007/s00221-022-06449-w

